# SARS-CoV-2 Omicron BA.2.75 variant may be much more infective than preexisting variants

**DOI:** 10.1101/2022.08.25.505217

**Authors:** Aki Sugano, Yutaka Takaoka, Haruyuki Kataguchi, Minoru Kumaoka, Mika Ohta, Shigemi Kimura, Masatake Araki, Yoshitomo Morinaga, Yoshihiro Yamamoto

**Affiliations:** Center for Clinical Research, Toyama University Hospital, Toyama 930-0194, Japan; Department of Medical Systems, Kobe University Graduate School of Medicine, Kobe, Hyogo 650-0017, Japan; Department of Computational Drug Design and Mathematical Medicine, Graduate School of Medicine and Pharmaceutical Sciences, University of Toyama, Toyama 930-0194, Japan; Data Science Center for Medicine and Hospital Management, Toyama University Hospital, Toyama 930-0194, Japan; Center for Advanced Antibody Drug Development, University of Toyama, Toyama 930-0194, Japan; Life Science Institute, Kobe Tokiwa University, Kobe, Hyogo 653-0838, Japan; Division of Genomics, Institute of Resource Development and Analysis, Kumamoto University, Kumamoto 860-0811, Japan; Department of Microbiology, Toyama University Graduate School of Medicine and Pharmaceutical Sciences, University of Toyama, Toyama 930-0194, Japan; Department of Clinical Infectious Diseases, Toyama University Graduate School of Medicine and Pharmaceutical Sciences, University of Toyama, Toyama 930-0194, Japan

**Keywords:** SARS-CoV-2, COVID-19, Infectivity, Spike protein, Docking affinity

## Abstract

**Objectives:** In our previous research, we developed a mathematical model via molecular simulation analysis to predict the infectivity of seven SARS-CoV-2 variants. In this report, we aimed to predict the relative risk of the recent new variants of SARS-CoV-2 as based on our previous research.

**Methods:** We subjected Omicron BA.4/5 and BA.2.75 variants of SARS-CoV-2 to the analysis to determine the absolute evolutionary distance of the spike protein gene (S gene) of the variants from the Wuhan variant so as to appreciate the changes in the spike protein. We performed the molecular docking simulation analyses of the spike proteins with human angiotensin-converting enzyme 2 (ACE2) to understand the docking affinities of these variants. We then compared the evolutionary distances and the docking affinities of these variants with those of the seven variants that we had analyzed in our previous research.

**Results:** The evolutionary distances of the S gene in BA.4/5 and BA.2.75 from the Wuhan variant were longer than those of other variants. BA.2.75 had the highest docking affinity of the spike protein with ACE2 (ratio per Wuhan variant).

**Conclusion:** The important results from this analysis are the following: BA.2.75 has both the highest docking affinity and the longest evolutionary distance of the S gene. These results suggest that BA.2.75 infection can spread farther than can infections of preexisting variants.

## 1. Introduction

In Japan, as of July 2022, infection with the Omicron BA.5 variant of SARS-CoV-2 has become an epidemic disease. In addition, Omicron BA.2.75 was discovered and is thought to present a particular risk inasmuch as it may cause a coming epidemic. We previously constructed a mathematical model to predict the infectivity of SARS-CoV-2 variants—Alpha, Beta, Gamma, Delta, Omicron BA.1and BA.2 as ratio per Wuhan variant (Takaoka et al., 2022). In this research here, we report the predicted risks for Omicron BA.4/5, and BA.2.75 which were recently recognized as being causes of epidemic diseases. For this purpose, we utilized the analyses of the evolutionary distance and the docking simulation that we established in our previous research (Takaoka et al., 2022).

## 2. Materials and methods

### 2.1 Determination of the absolute evolutionary distances between the Wuhan variant and variant spike protein genes (S genes), and docking affinities of the different spike proteins with ACE2

We analyzed the absolute evolutionary distances of the S gene from the Wuhan variant for the variants that we used—Alpha, Beta, Gamma, Delta, Omicron BA.1, BA.2, BA.4/5, and BA.2.75 via the ClustalW program (Thompson et al., 1994) and FastTree program (Price et al., 2009). We obtained the sequences of the S gene by searching NCBI (MN908947 for Wuhan, OW519813 for Alpha, OM791325 for BA.1) (National Library of Medicine) or the EpiCoV database of GISAID for the complete sequence of the S gene (EPI_ISL_5142896 for Beta, EPI_ISL_14534452 for Gamma, EPI_ISL_4572746 for Delta, EPI_ISL_13580480 for BA.2, EPI_ISL_13304903 for BA.4/5, and EPI_ISL_14572678 for BA.2.75) (Elbe et al., 2017; Khare et al., 2021).

We used docking simulation and cluster analysis to investigate the docking affinity of the receptor binding domain (RBD) of each variant spike protein with ACE2 (Takaoka et al., 2022). We calculated the docking affinity by means of the following expression:

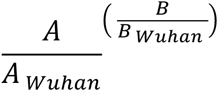

where A is the average score of the most stable cluster and B is the score of the most stable complex in the docking result.

We obtained the amino acid sequences of the spike proteins from the CoVariants website, which classifies the variants according to Nextstrain clades: 22A (Omicron, BA.4), 22B (Omicron, BA.5), and 22D (Omicron, BA.2.75) (Hadfield et al., 2018; Hodcroft 2021). BA.4 and BA.5 have the identical spike proteins, and thus we grouped the sequence of their S genes or their amino acid sequences together in this research.

## 3. Results

### 3.1. Absolute evolutionary distances for S gene variants and results of docking of the RBD with ACE2 protein

Table 1 shows the absolute evolutionary distances of the S gene between the Wuhan variant and each of the other variants, as well as the docking affinities of the RBD of the spike protein with ACE2 (ratio per Wuhan variant), which we determined from the docking simulation. The variants with longer evolutionary distances from the Wuhan variant had a tendency toward causing more epidemics. The Omicron BA.2.75 variant had the highest docking affinity of the spike protein with the ACE2 protein.

**Table 1.** The evolutionary distance of the S gene and the docking affinity of spike protein with ACE2 (ratio per Wuhan variant)

|  | Wuhan | Alpha | Beta | Gamma | Delta | Omicron<br>BA.1 | Omicron<br>BA.2 | Omicron<br>BA.4/5 | Omicron<br>BA.2.75 |
| --- | --- | --- | --- | --- | --- | --- | --- | --- | --- |
| Absolute values of<br>the Evolutionary<br>Distance of S gene<br>(from Wuhan) x 10<sup>-3</sup> | - | 2.07 | 2.07 | 3.55 | 3.26 | 10.72 | 8.31 | 9.21 | 10.99 |
| Docking affinity<br>(Ratio per Wuhan) | 1 | 1.15 | 1.23 | 1.32 | 2.40 | 3.00 | 7.91 | 4.82 | 21.05 |

In comparison with BA.2, BA.4/5 had a longer S gene evolutionary distance but showed a lower docking affinity. BA.2.75 had not only a longer evolutionary distance but also a much higher docking affinity than did BA.2.

## 4. Discussion

For this report, we analyzed the evolutionary distance of the S gene and the docking affinity with ACE2 of the recent new Omicron variants, BA.4/5 and BA.2.75. We focused on only the S gene sequence to calculate the absolute evolutionary distance and did not use the genome sequence, which are used for the phylogeny provided by Nextstrain (Hadfield et al., 2018).

Our analyses is based on the following two factors, which play important roles in the infectivity of the virus variants: (1) the ability of the virus to enter human cells and (2) the effect of a neutralizing antibody in humans or the effect of vaccines. The first is demonstrated by the docking affinity of each RBD in the spike protein with ACE2, that is, which docking affinity is greater. The second is shown by the evolutionary distance of the S gene from the Wuhan variant: the longer the distance is, the weaker the effect of vaccines. It is because the currently available vaccines were developed on the basis of the Wuhan variant (Hassine 2022). However, the risk for exacerbation of SARS-CoV-2 cannot be appreciated via the methods for the two factors.

Omicron BA.2.75 showed the highest docking affinity of the spike protein with the ACE2 protein compared with the other seven variants (ratio per Wuhan variant). In addition, the S gene evolutionary distance of BA.2.75 from the Wuhan variant was the longest of the variants. Therefore, BA.2.75 has a superior ability to enter human cells, and also the current vaccines are much less effective against this variant. These results suggest that the BA.2.75 infection can spread farther than can infections of preexisting variants. In addition, our results indicate the need for a great caution in managing BA.2.75 because the number of severely ill patients or sufferers will be increased along with the increased number of infected individuals even if this variant has low risk for exacerbation.

## 5. Conclusion

We demonstrated here that the Omicron BA.2.75 variant of SARS-CoV-2 has the longest results with regard to the evolutionary distance of the S gene from the Wuhan variant and the highest results of the docking simulation for spike protein with ACE2. Our results indicate that Omicron BA.2.75 poses a greater risk to global health than other variants and that we must pay close attention to the Omicron BA.2.75 infection trends.

## Abbreviations

S gene: spike protein gene
ACE2: angiotensin-converting enzyme 2
RBD: receptor binding domain

## Data availability

Data that support the findings of this study are available from the corresponding author upon reasonable request, except publicly available data source.

## Acknowledgments

We gratefully acknowledge all data contributors, i.e., the Authors and their Originating laboratories responsible for obtaining the specimens, and their Submitting laboratories for generating the genetic sequence and metadata and sharing via the GISAID Initiative and all data provided by CoVariants, on which this research is based.

## Author contributions

Y.T. conceived and designed this research. Y.T. and A.S. preformed the analyses and acquired the data. Y.T., A.S., and M.O. interpreted the data. Y.T. and A.S. wrote the draft, and all authors reviewed and approved the manuscript.

## Ethical approval statement

This research is not applicable because we performed computer analyses by using sequence data obtained from public database.

